# Effects of sugars and lipids on the growth and development of ***Caenorhabditis elegans***

**DOI:** 10.1101/615807

**Authors:** Xiong Wang, Lin Zhang, Lei Zhang, Wenli Wang, Sihan Wei, Jie Wang, Huilian Che, Yali Zhang

## Abstract

Excessive intake of carbohydrates and fats causes over-nutrition, leading to a variety of diseases and complications. Here, we characterized the effects of different types of sugar and lipids on the growth and development of *Caenorhabditis elegans*. We measured the lifespan, reproductive capacity, and length of nematodes after sugars and lipids treatment alone and co-treatment of sugars and lipids. Furthermore, by using transcriptome sequencing technology, we studied the mechanisms underlying the damaged caused by high-sucrose and high-stearic acid on *C. elegans*. The results showed that a certain concentration of sugar and lipid promoted the growth and development of nematodes. However, excessive sugars and lipids shortened the lifespan and length of nematodes and destroyed their reproductive capacity. Based on the results of the orthogonal test, we selected 400 mmol/L sucrose and 500 μg/mL stearic acid to model a high-sugar and high-lipid diet for *C. elegans*. High-sugar and high-lipid intake altered the expression of genes involved in biofilm synthesis, genes that catalyze the synthesis and degradation of endogenous substances, and genes involved in innate immunity, resulting in physiological damage.

## Introduction

All animals require energy to sustain basic life activities, such as survival, growth, and reproduction. Digested and absorbed, dietary nutrients are important precursors for the synthesis and metabolism of cells. Carbohydrates and fats are the main organic material sources to sustain life activities. Carbohydrates are present in all living organisms and have a variety of basic functions, providing energy for all non-photosynthetic organisms. Lipids perform many essential functions in cells. Due to their highly reduced state, they are effective energy storage molecules. They are bilayered hydrophobic units that form cells and organelle membranes, and act as effective signaling molecules to facilitate communication between cells (Satouchi *et al*. 1993). Reasonable carbohydrate and fat intake has a positive impact on human life activities, but excessive intake may be harmful to the human body, leading to diabetes, high blood pressure, and tumors. In recent years, with the prevalence of human obesity and diabetes, interest in lipid and carbohydrate metabolism has become increasingly prominent.

With the improvement in people’s living standards, the dietary structure has gradually developed towards high sugar and high lipid. Continued high sugar and high lipid intake can lead to several abnormal conditions, such as obesity and type 2 diabetes. High fat and high lipid intake lead to over-nutrition, which, in turn, causes obesity. In the past 40 years, the world’s obese population has increased from 105 million in 1975 to 641 million in 2014. Almost one in every 8 adults in the world has obesity problems; China is a country with the greatest number of obese people (Ashrafi 2007). Obesity increases the risk of type 2 diabetes, cardiovascular disease, stroke, high blood pressure, and cancers, affecting physical health. Studies have found that obesity is affected by age, diet, living environment, and genes (Ezzati *et al*. 2017). Obesity is essentially an energy balance disorder caused by excessive energy intake over energy consumption (Collaboration 2016). Energy balance is highly regulated and complexly related to energy consumption by food sensory, nutrient intake signals, nutrient delivery and storage, eating behavior, growth, reproduction, basal metabolism, and physical activity. The integrated metabolic system inside the human body is highly complex and redundant, and it is difficult to fully elucidate the mechanisms underlying human obesity in a short period of time (Badman and Flier 2007). In addition, mammalian genetic experiments take a long period of time. Therefore, many researchers are trying to study obesity-related metabolism in lower model organisms. *Caenorhabditis elegans* has been widely used to study obesity-related metabolism due to several advantages: (1) knowledge of the complete genome sequence; (2) the core genes involved in lipid and sugar metabolism pathways are highly conserved and align with higher organisms; and (3) low price, short life cycle, operability, transparent, and easy to observe.

Resveratrol is a natural plant antibody produced when plants encounter external stimuli, such as fungi and ultraviolet radiation, and plays important role in protecting plants (Pangeni *et al*. 2014). Resveratrol has been derived from various parts of several plants, including the fruits, skin, and seeds. Numerous studies have shown that resveratrol exhibits various biological activities, such as blood fat-lowering, antioxidative, anti-aging, anti-tumor, anti-thrombosis, and immunoregulatory effects (Aschemann-Witzel and Grunert 2015). In terms of lipid metabolism, resveratrol inhibits fat accumulation by reducing the synthesis of lipids and cholesterol, while promoting fat decomposition by enhancing fatty acid oxidation and glucose transport (GÓmezzorita *et al*. 2012). Resveratrol ameliorates the abnormal lipid metabolism induced by dietary fat. The greater the concentration of resveratrol within a certain range, the better the recovery of antioxidant capacity in mice and the better the ability to improve lipid metabolism. However, after a certain range, resveratrol causes pro-oxidation in the body, and does not improve hepatic redox status and lipid metabolism(Ran *et al*. 2017).

In the present study, we evaluated the effects of sugars and lipids on the damage caused in C. elegans and selected the appropriate sugar and lipid concentration to model a high-sugar and high-fat diet. In addition, we explored the role of resveratrol in protecting C. elegans against high-sugar and high-lipid damage. Moreover, by using transcriptome sequencing technology, we studied the damage mechanism of high sucrose and high stearic acid on C. elegans and the repair mechanism of resveratrol.

## Materials and methods

### Material

Resveratrol (99%) was purchased from Sigma (Sigma, America). The sucrose, fructose, glucose, stearic acid, cholesterol, and linoleic acid used in the tests were of analytical grade and purchased from Sigma (Sigma, America). Stock solutions (200 mM) of resveratrol in dimethyl sulfoxide (DMSO) were stored at −20 °C.

### Animals, culture, and treatment with resveratrol

Wild type N2 strains were obtained from the Caenorhabditis Genetics Center and maintained on nematode growth medium (NGM) with concentrated Escherichia coli OP50 as a food resource, at 20 ℃. Age-synchronized worms were generated in all experiments using the sodium hypochlorite method. Resveratrol was dissolved in DMSO to a final concentration of 50 mg/mL and added at an appropriate ratio to molten agar NGM.

### Life span

Life span analyses were performed as previous described, at 20 ℃ (Lee *et al*. 2009). L1 larvae were placed onto a sugar-containing NGM plate or a lipid-containing NGM plate, and then, the live nematodes in the plate were transferred to a fresh plate every day. The number of nematodes surviving was recorded each day until all died. The death of nematodes was defined as no reflection when gently prodded with a platinum wire. Lost nematodes and dead nematodes as they climb to the wall of the culture medium should were excluded from the statistics. Each experimental group consisted of 10 nematodes.

### Reproduction capacity

Reproduction capacity was analyzed as previously described. L4 larvae from the synchronized L1 generation were placed onto an individual NGM plate. Nematodes were transferred to a new medium every day until end of reproduction. Approximately after 12 hours, the number of eggs on the old medium was counted. Finally, the total amount of eggs laid by nematodes in the whole life was counted. Each experimental group consisted of 10 nematodes.

### Measurements of body length

Animals were grown at 20 ℃. After treatment of the sample, the synchronic larvae were picked from the NGM culture plate and placed under a stereo microscope. The culture dish was rotated to make the body of the nematode closer to the scale and the length of the body was evaluated. According to the ratio of the scale to the actual length, the body length of the nematode was calculated. The length of the nematode was measured and recorded every 24 hours until the sixth day. Each experimental group consisted of 10 nematodes.

### Total RNA extraction, library preparation, and RNA-seq

Trizol method was used to extract total RNA from nematodes, including control group (control), after sucrose treatment at concentration of 400 mmol/L (suc), stearic acid treatment at concentration of 500 μg/mL (ste), co-treatment with 400 mmol/L concentration of sucrose and 500 μg/mL concentration of stearic acid (suc-ste), and co-treatment with sucrose-stearic acid-500 μg/mL resveratrol (suc-ste-res). Each group was analyzed in triplicates. Total RNA was quantified using Nanodrop spectrophotometer. The RNA of each sample that passed quality control test was used for library construction. The cDNA library construction and sequencing on Illumina Hiseq X Ten were performed at Beijing Mega Genomic Technology (Beijing, China), following the manufacture’s standard protocol.

### Analysis of RNA-seq

By filtering rRNA reads, sequencing adapters, short-fragment reads, and other low-quality reads, clean reads were obtained. The clean reads were mapped to nematodes reference genome (National Center Biotechnology Information reference sequence: GCF_000002985.6) by Tophat v2.1.0.

In order to assess the quality of the sequencing, gene coverage and sequencing saturation were analyzed. After genome mapping, the open source suite of the tool Cuffinks was run with a reference annotation to generate fragments per kilo base of exon per million mapped read (FPKM) values for standardized calculation of the gene-expression levels. Differentially expressed genes (DEGs) were identified using Cuffdiff software. The calculated gene expression levels could thus be used for comparing gene expression directly between the different samples. The significance threshold of the p-value of multiple tests was set by the false discovery rate (FDR). Fold-change in expression was also estimated according to the FPKM in each sample. Differentially expressed genes were selected using the following filter criteria: FDR ≤ 0.05 and fold-change ≥ 2.

The DEGs were subjected to enrichment analysis of Gene Ontology (GO) and Kyoto Encyclopedias of Genes and Genomes (KEGG). GO functions and KEGG pathways were analyzed by Blast2GO software (https://www.blast2go.com/) and Blastall software (http://www.kegg.jp/).

### Statistical analyses

Results are expressed as mean ± SEM. Statistical significance was determined using one-way analysis of variance (ANOVA) followed by Tukey’s multiple-comparison test with SPSS version 19.0. Differences were considered significant when p < 0.05.

### Data Availability

Upon request, all the data will be available to journal and readers. Seven supplementary figures and three tables are available at Figshare.

## Results

### Effect of sugar and lipids on lifespan of N2

Nematodes were treated with sugar at concentrations ranging from 0 to 550 mmol/L and lipid at concentrations ranging from 0 to 600 μg/mL. As shown in Table 1, the average lifespan of nematodes treated with different concentrations of sucrose, fructose, and glucose increased initially, and then decreased, which was consistent with previous reports (Schlotterer *et al*. 2009; Zheng *et al*. 2017). Treatment with sucrose and fructose at a concentration of 5 mmol/L had a weak effect on the average lifespan of nematodes, whereas treatment with 5 mmol/L glucose significantly prolonged the average lifespan of nematodes. Treatment with 50 mmol/L sucrose, fructose, and glucose significantly prolonged the average lifespan of nematodes and delayed the onset of death. Treatment with sucrose at concentrations above 400 mmol/L significantly shortened the average lifespan of nematodes, whereas for fructose and glucose, the tuning points were 500 mmol/L and 520 mmol/L, respectively. This indicates that treatment with low concentrations of sucrose, fructose, and glucose prolonged the average lifespan of nematodes, whereas when the concentration of sugar reached a certain level, the average lifespan of nematodes was significantly shortened. Among the three kinds of sugar, sucrose exhibited a relatively narrow range of concentration that prolonged the lifespan of nematodes, but glucose had a wider range of said concentration—5 mmol/L to 500 mmol/L.

**Table 1.**
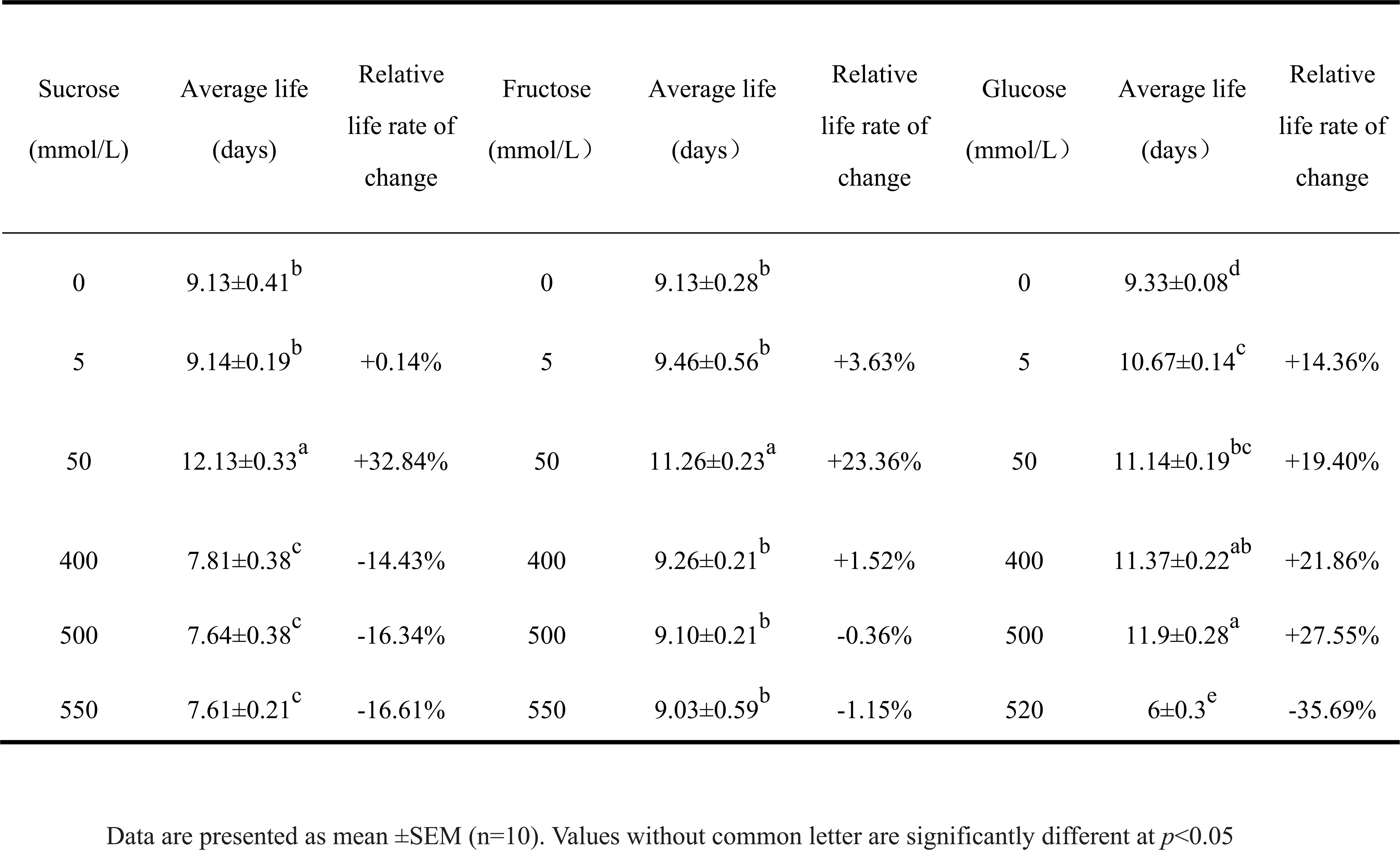
The mean life span of N2 in different sugars

As shown in Table 2, the average lifespan of nematodes treated with different concentrations of stearic acid, linoleic acid, and cholesterol increased initially, and then decreased. Treatment with 5 μg/mL of stearic acid and linoleic acid had weak effect on the average lifespan of nematodes, whereas treatment with 5 μg/mL of cholesterol significantly prolonged the average lifespan of nematodes. This was consistent with a previous report (Dou *et al*. 2012). Treatment with 50 μg/mL and 200 μg/mL of stearic acid and linoleic acid significantly prolonged the average lifespan of nematodes. As expected, high concentrations of lipid began to shorten the lifespan of nematodes. Stearic acid can prolong the average life span of nematodes by up to 31.82% at a concentration of 50 μg/mL, but it decreases the average life span severely at a concentration of 600 μg/mL. Although linoleic acid also shows a similar pattern as the other test substances, it did not decrease the average life span of nematodes at any higher concentration we used in the experiments compared to that of controls. As *C. elegans* cannot synthesize cholesterol itself, 5 μg/mL of cholesterol was added to the control medium in every experiment except in the cholesterol test, in which no cholesterol was added to the control medium. Our result showed that 5 μg/mL of cholesterol is the best concentration to prolong the average life span of nematodes.

**Table 2.**
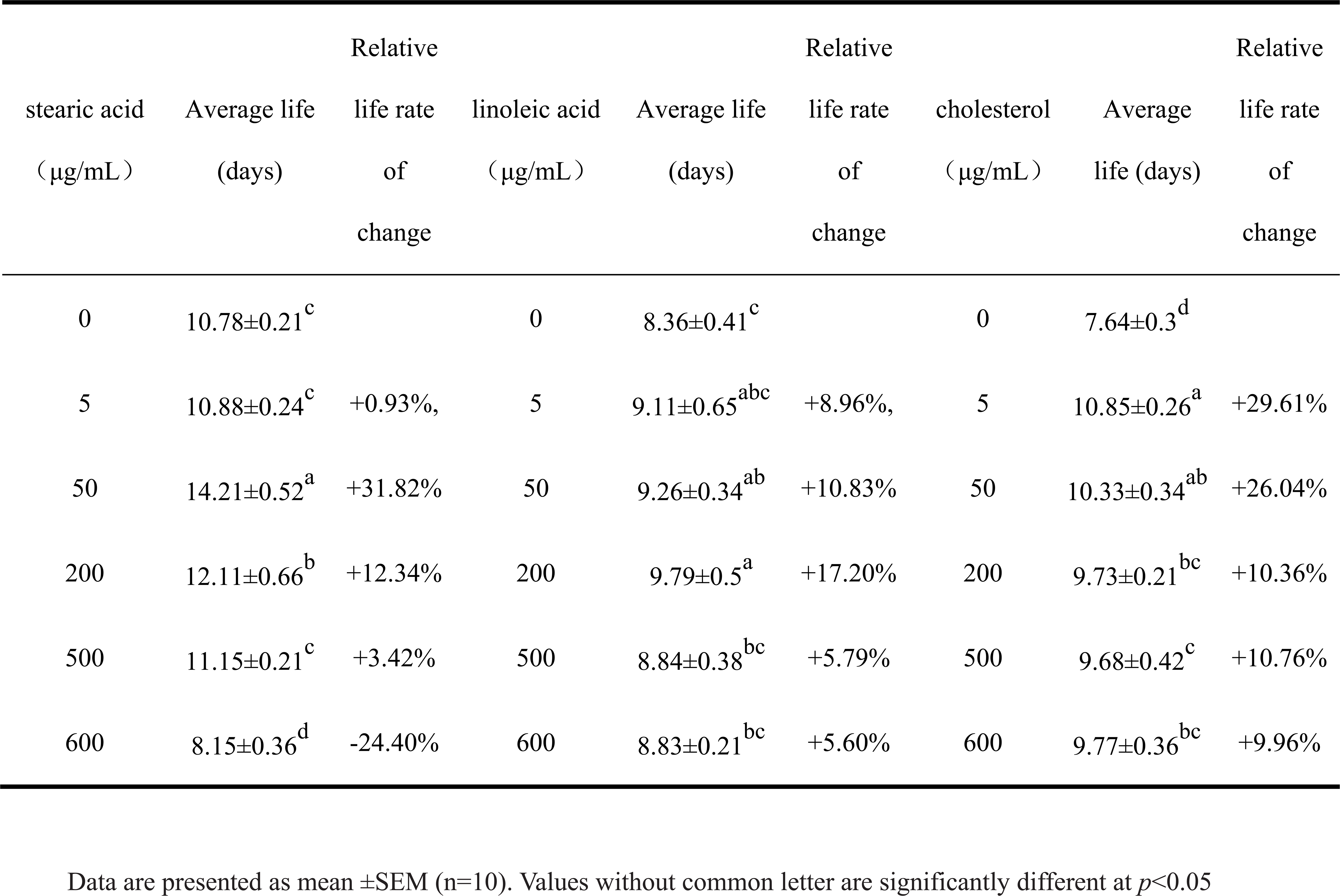
The mean life span of N2 in different lipids

### Effect of sugar and lipids on the reproductive capacity of N2

As shown in Fig. 1A, after treatment with sucrose, fructose, and glucose, the total number of eggs laid by nematodes increased initially, and then decreased along with the increase in sucrose concentration. After treatment with 400, 500, and 550 mmol/L sucrose, the number of eggs decreased by 61.57%. 65.97%, and 79.1%, respectively (Fig.1B). Under the treatment with low concentration of fructose, the number of eggs laid by nematodes increased with the increase in fructose concentration. However, treatment with fructose above 400 mmol/L significantly reduced the number of eggs laid by nematodes (Fig.1C). Treatment with 0 to 50 mmol/L glucose had no effect on the egg production of nematodes. After treatment with 400, 500, and 520 mmol/L glucose, the total number of eggs laid by nematodes decreased by 36.92%, 71.62%, and 86.98%, respectively (Fig.1D). Taken together, high concentration sugar intake exhibited significant damage to the reproductive capacity of nematodes, and the damage increased with increasing concentration. After reaching a certain level, the nematode eventually loses its reproductive ability. As described in Fig1B-D, the nematodes treated with control and low concentration of sugar entered the spawning period on the third day and ended the spawning on the sixth day. For concentration higher than 400 mmol/L sugar group, the spawning periods were delayed 1-2 days and some lasted one day more (from the 4^th^-5^th^ day to 7^th^ −8^th^ day), except for 520 mmol/L glucose treated group, which started to lay eggs on 8^th^ day and ended on 12^th^ day. Delay of spawning period means inhibition of nematodes’ development, which occurs most severely in 520 mmol/L glucose treated group. Moreover, the higher the concentration of sugar is, the less eggs the nematodes lay and the further beginning day of laying eggs.

**Figure 1.**
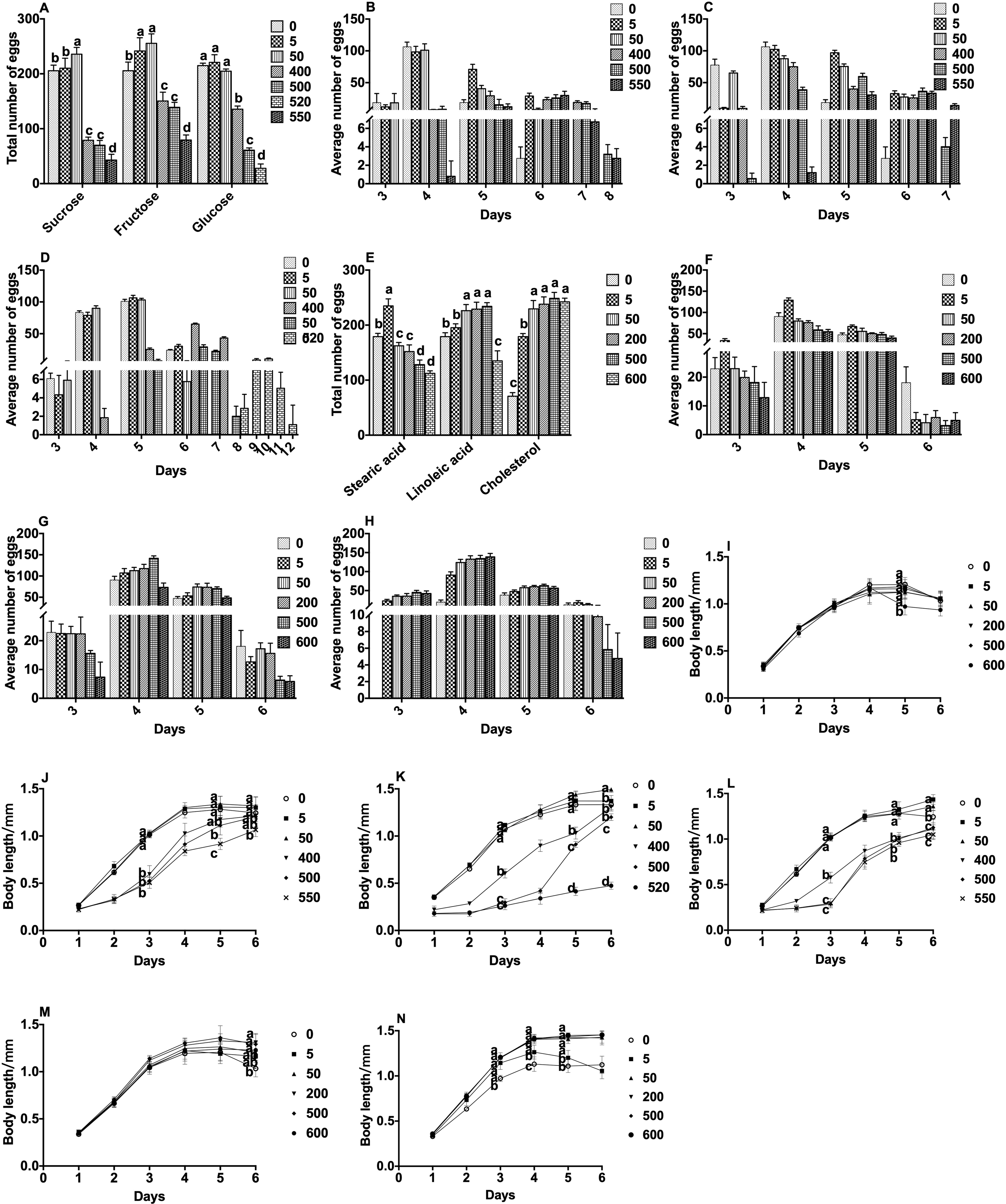
Effects of sugars and lipids on reproduction and body length of nematode. **A**, the total number of offspring in the entire spawning period of the nematode under each concentration gradient of sugars. **B-D**, the effects of sucrose(**B**), fructose (**C**) and glucose **(D**) on the lifespan of nematodes. **E**, the total number of offspring in the entire spawning period of the nematode under each concentration gradient of lipids. **F-H**, the effects of stearic acid (**F**), linoleic acid (**G**) and cholesterol (**H**) on the reproductive capacity of nematodes. **I-K**, the effects of sucrose (**I**), fructose (**J**) and glucose (**K**) on the body length of nematodes. **L-N**, the effects of stearic acid (**L**), linoleic acid (**M**) and cholesterol (**N**) on the body length of nematodes. Data are presented as mean ±SEM (n=10). Values without common letter are significantly different at p<0.05.

As shown in Fig.1E, after treatment with stearic acid and linoleic acid, the total number of eggs laid by nematodes increased initially, and then, decreased along with the increase in stearic acid and linoleic acid concentration. Interestingly, cholesterol treatment at low concentrations significantly increased nematode spawning. When the cholesterol concentration was greater than 50 μg/mL, there was no significant difference in the amount of eggs laid by nematodes at any concentration. This result was similar to the lifespan of nematodes, indicating that when cholesterol is added above 50 μg/mL, the nematode’s demand for cholesterol is saturated. In addition, different lipid treatments have different turning points in reducing the levels of nematode spawning. The treatment with stearic acid at a concentration of 50 μg/mL reduced the number of eggs laid, whereas linoleic acid at a concentration of 600 μg/mL only destroyed the nematode reproductive ability (Fig.1F-H). The greater the concentration of stearic acid, the stronger was the damage. The sperm plasma membrane of nematodes is rich in cholesterol, and the survival of sperm requires the supply of exogenous cholesterol (Matyash *et al*. 2001; Dou *et al*. 2012). Since the nematode does not synthesize cholesterol itself, the total amount of progeny of the nematode after treatment with cholesterol increases initially, and then, decreases slightly with the increase in cholesterol concentration. Shim Y H et al (Shim *et al*. 2002) reported a significant decrease in the number of eggs laid by nematodes and a decrease in growth rate after blocking the supply of exogenous cholesterol. This is consistent with our findings that a certain concentration of cholesterol increases the reproductive capacity of nematodes. In general, the effect of lipids on the spawning of nematodes was not as severe as the effect of sugar.

### Effect of sugar and lipids on the body length of N2

Nematodes need to consume energy for their growth and spawning. Sugar, as a nutrient, can provide a lot of energy for the life activities of nematodes. As shown in Fig.1I, the body length of the nematodes treated with 5 mmol/L and 50 mmol/L sucrose was similar to the body length of the nematodes in the control group. After the nematodes entered the spawning period, they continued to grow and the body length was about 1.1 times longer than that in the control group organisms. This indicated that the sucrose concentration in the range of 5 to 50 mmol/L did not change the length of the nematodes; however, it can promote the growth of nematodes during the spawning period and increase the maximum length of the nematodes. In addition, high concentrations of sucrose shorten nematode length. Treatment with fructose at concentrations of 5 to 50 mmol/L had no effect on the length of the nematode, but higher concentrations of fructose significantly shortened nematode’s maximum length (Fig.1J). Treatment with 5 mmol/L glucose had no effect on the length of the nematode. During the spawning period, treatment with 50 mmol/L glucose promoted the growth of the nematode and increased the length of the nematode (Fig.1K). Treatment with higher concentrations of glucose significantly shortened nematode length, especially for 520 mmol/L glucose treated group, which only grew up to 1/3 length of control nematodes on 6^th^ day.

As shown in Fig.1L-N, compared with the length in the control group, except for treatment with stearic acid at a concentration of 600 μg/mL, there was no significant difference in the length of nematodes after treatment with the other concentrations of stearic acid. In the growing season, the body length of the nematode grew rapidly and reached a maximum of 1.2 mm on the 4^th^ day, after treatment with stearic and linoleic acid (Fig.1L&M). Treatment with a concentration of linoleic acid above 200 μg/mL delayed the appearance of nematode aging but did not change the maximum length of the nematode. Previous studies have found that body length of nematode grown on cholesterol-added media is longer than that grown on cholesterol-free medium (Shanmugam *et al*. 2017). In our study, after treatment with cholesterol at the concentration of 50 μg/mL, 200 μg/mL, 500 μg/mL, and 600 μg/mL, the growth rate of nematodes was basically the same as that of the control group during the growth phase. The body length of nematodes after cholesterol treatment showed significant difference from the 3^rd^ day, and reached the maximum length of 1.4 mm on the 4^th^ day, which was 1.1 times the length of the nematode in the control group.

### Effects of various levels of sucrose and stearic acid orthogonal design on the lifespan of N2

The lifespan of nematodes after treatment with different concentrations of sucrose and stearic acid is shown in Table 3. In the case of lower sugar concentrations of 0 to 250 mmol/L, it can be seen that the lifespan of nematode only treated with 50 μg/mL stearic acid was significantly prolonged. However, at a sugar concentration of 400 mmol/L, an increase in the concentration of stearic acid exhibited a tendency to shorten the lifespan of nematodes. There was no significant difference in the lifespan of nematodes treated with different concentrations of stearic acid at constant sucrose concentration of 400 mmol/L. In addition, in the case of treatment with constant stearic acid concentration, the lifespan of the nematode increased initially, and then, decreased with the increase in sucrose concentration. This is consistent with the previous single sucrose treatment results. We observed that co-treatment with low concentration of sugar and lipid exhibited synergistic effect of extending the lifespan of nematodes. For example, after 50 mmol/L of sugar and 50 μg/mL of stearic acid co-treatment, the average lifespan of nematodes reached a maximum of 12.96 days, and the relative average life change rate was 31.25%.

**Table 3.**
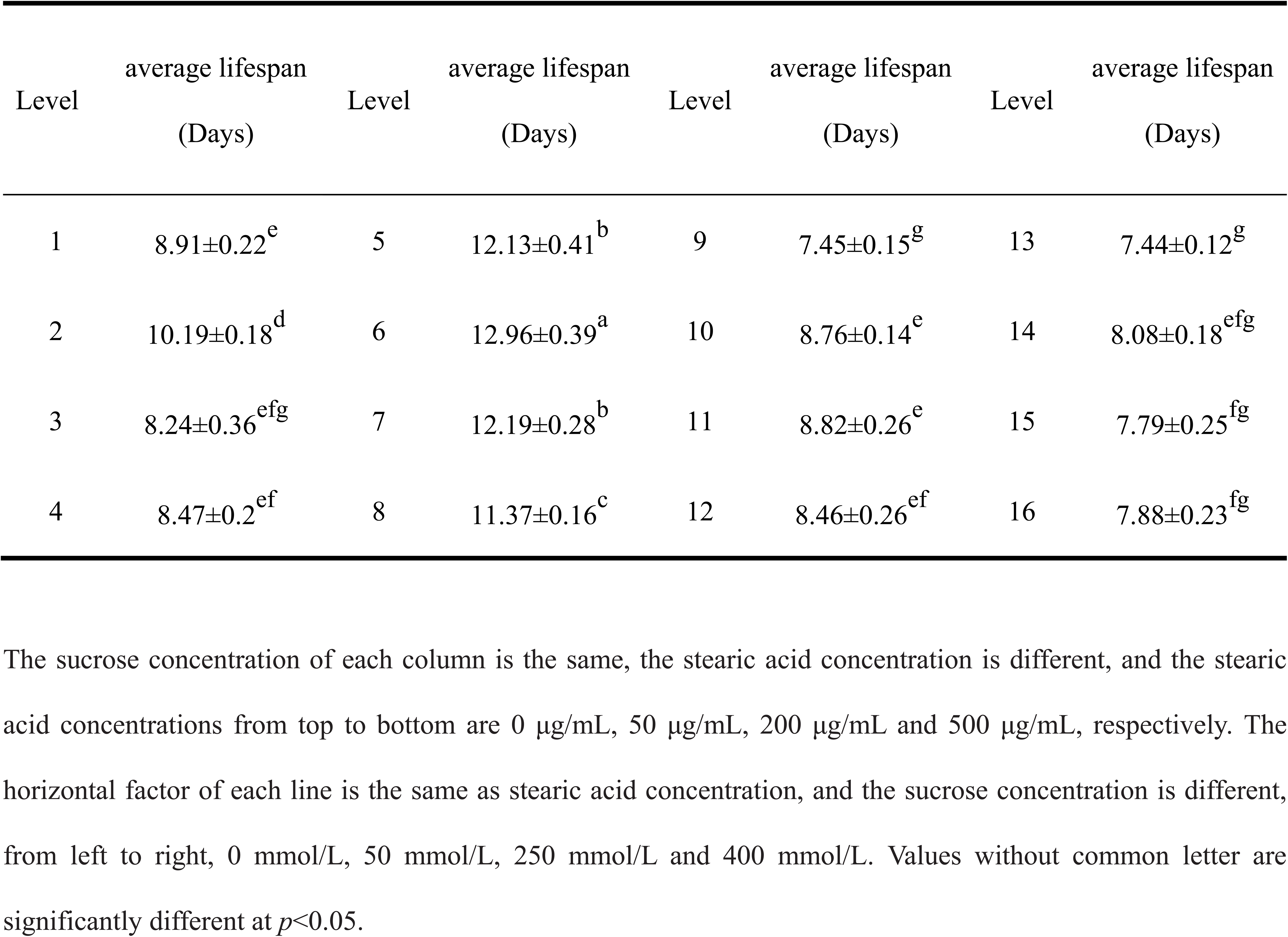
The effects of various levels of sucrose and stearic acid orthogonal design on the lifespan of nematodes.

### Effects of various levels of sucrose and stearic acid orthogonal design on the reproductive capacity of N2

As shown in Fig.2A, under the constant concentration of stearic acid, the total number of nematode offspring increased initially, and then, decreased with the increase of sucrose concentration, and reached a maximum at a concentration of 50 mmol/L sucrose. This was similar to the result of treatment of nematodes with sucrose alone. Under the constant sucrose concentration, the total number of nematode offspring gradually decreased with the increase in stearic acid concentration. This result is also in accord with the former result, in which the total number of eggs of nematodes began to decrease at concentration higher than 50 μg/mL (Fig.1A). The decrease in number of eggs after treatment with stearic acid began at a lower concentration than that of lifespan, and it kept such tendency even at different sucrose concentrations. In high sucrose concentration group, stearic acid and sucrose exhibited synergistic effect on decrease in number of eggs. When comparing the number of nematode offspring at each level of treatment, we found that the total number of nematode offspring in group co-treated with 400 mmol/L sucrose and 500 μg/mL stearic acid was the lowest.

**Figure 2.**
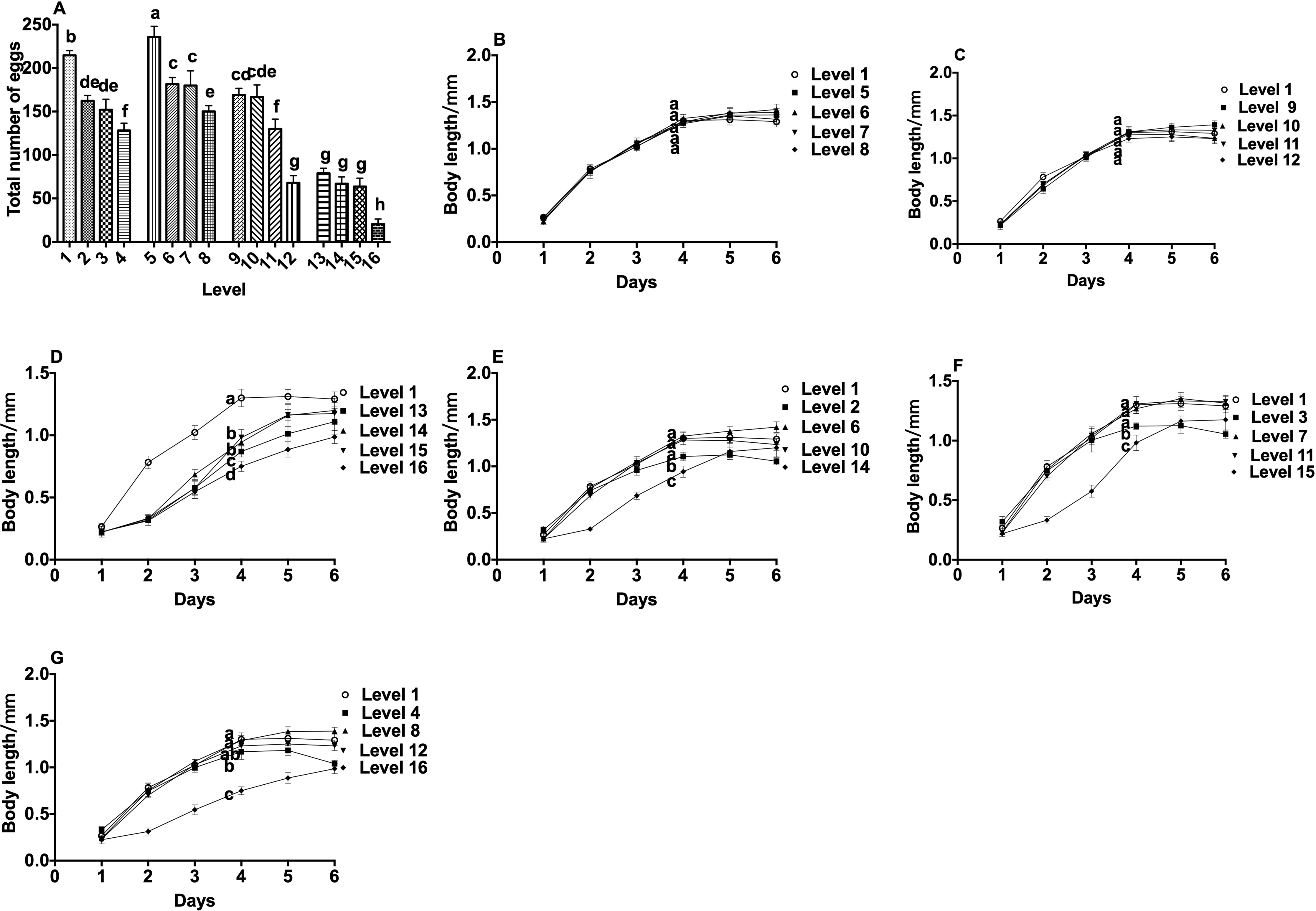
Effects of various levels of sucrose and stearic acid orthogonal design on reproduction and body length of nematode. **A**, The effects of various levels of sucrose and stearic acid orthogonal design on the reproductive capacity of nematodes. **B-G**, the effects of various levels of sucrose and stearic acid orthogonal design on the body length of nematodes. **B-D**: The same concentration of sucrose (50mmol/L, 250mmol/L, 500mmol/L, respectively), different concentration of stearic acid (0μg/mL, 50μg/mL, 200μg/mL and 500μg/mL, respectively) in the same figure. **E-G**: The same concentration of stearic acid (50μg/mL, 200μg/mL, 500μg/mL, respectively), different concentration of sucrose (0mmol/L, 50mmol/L, 250mmol/L and 400mmol/L, respectively) in the same figure. Data are presented as mean ±SEM (n=10). Values without common letter are significantly different at p<0.05.

### Effects of various levels of sucrose and stearic acid orthogonal design on the body length of N2

As shown in Fig.2B-D, at 50 mmol/L and 250 mmol/L sucrose concentration, there was no significant difference in the length of nematodes treated with different concentrations of stearic acid. The maximum length of nematode was 1.42 ± 0.083 mm and 1.39 ± 0.083 mm, respectively. At a concentration of 400 mmol/L sucrose, treatment with stearic acid at concentrations of 50 μg/mL and 200 μg/mL significantly increased nematode length, reaching 14% and 16%, respectively. In addition, treatment with 500 µg/mL stearic acid significantly inhibited nematode growth, and the maximum length of nematodes was 1.19 ± 0.088 mm (Fig.2D). Under treatment with the same concentration of stearic acid, treatment with low concentration of sucrose had no effect on nematode length, but high concentration of sucrose significantly shortened nematode length. In addition, at 50 μg/mL and 200 μg/mL stearic acid concentrations, treatment with 250 mmol/L sucrose increased the maximum length of nematode, increasing by 14.02% and 20.15%, respectively (Fig.2E&F). However, at any concentration of stearic acid, treatment with sucrose at concentration of 400 mmol/L significantly shortened the length of the nematode.

Thus, we observed that sucrose has a more dramatic effect on nematode lifespan, growth, and reproduction. Low concentration of sucrose (50 mmol/L) had no significant effect on the growth and development of nematodes, but it significantly promoted the body length of adult nematodes. At the same time, it significantly increased the number of eggs laid by nematodes and significantly prolongs the lifespan of nematodes. Medium concentration of sucrose (250 mmol/L) also promoted the growth of nematode adults, but has no significant effect on the number of eggs. However, high concentrations of sucrose (400 mmol/L) significantly reduced the number of eggs and shortened the lifespan of nematodes.

The effect of stearic acid on nematodes is less prominent than that of sucrose. It also prolonged the lifespan of nematodes at low concentrations (50 μg/mL) and worked synergistically with 50 mmol/L sucrose. Moreover, it showed inhibition of nematode reproduction ability at each gradient sucrose concentration. Furthermore, its effect on the growth and development of nematodes and adult body length was not significant. The decrease in the lifespan of nematodes after treatment with high concentration of stearic acid (400 μg/mL) was much less than that after sucrose treatment (4.94% vs. 16.5%). However, when it is co-treated with sucrose, the growth and development of nematodes, the length of adult worms, and the number of eggs laid, are more significantly inhibited.

### Effect of resveratrol on lifespan, reproductive ability, and body length of N2

Recently, the anti-aging effect of resveratrol has increasingly gained attention. In our experiment, we explored the repair effect of resveratrol on sucrose-stearic acid damage to nematodes. As shown in Table 4, there was no significant difference in the average lifespan of nematodes between the 50 μg/mL and 100 μg/mL resveratrol treated groups, compared to that in the control group. This indicated that resveratrol, at concentrations below 100 μg/mL, exhibited less prominent effect on the average lifespan of nematodes treated with sucrose-stearic acid. However, after treatment with resveratrol at the concentration of 500 μg/mL, 750 μg/mL, and 1000 μg/mL, the lifespan of nematodes was significantly prolonged in a dose-dependent manner. Our results were consistent with the previously reported results (Desjardins *et al*. 2017). However, there was no significant difference between the groups treated with different resveratrol concentrations.

**Table 4.**
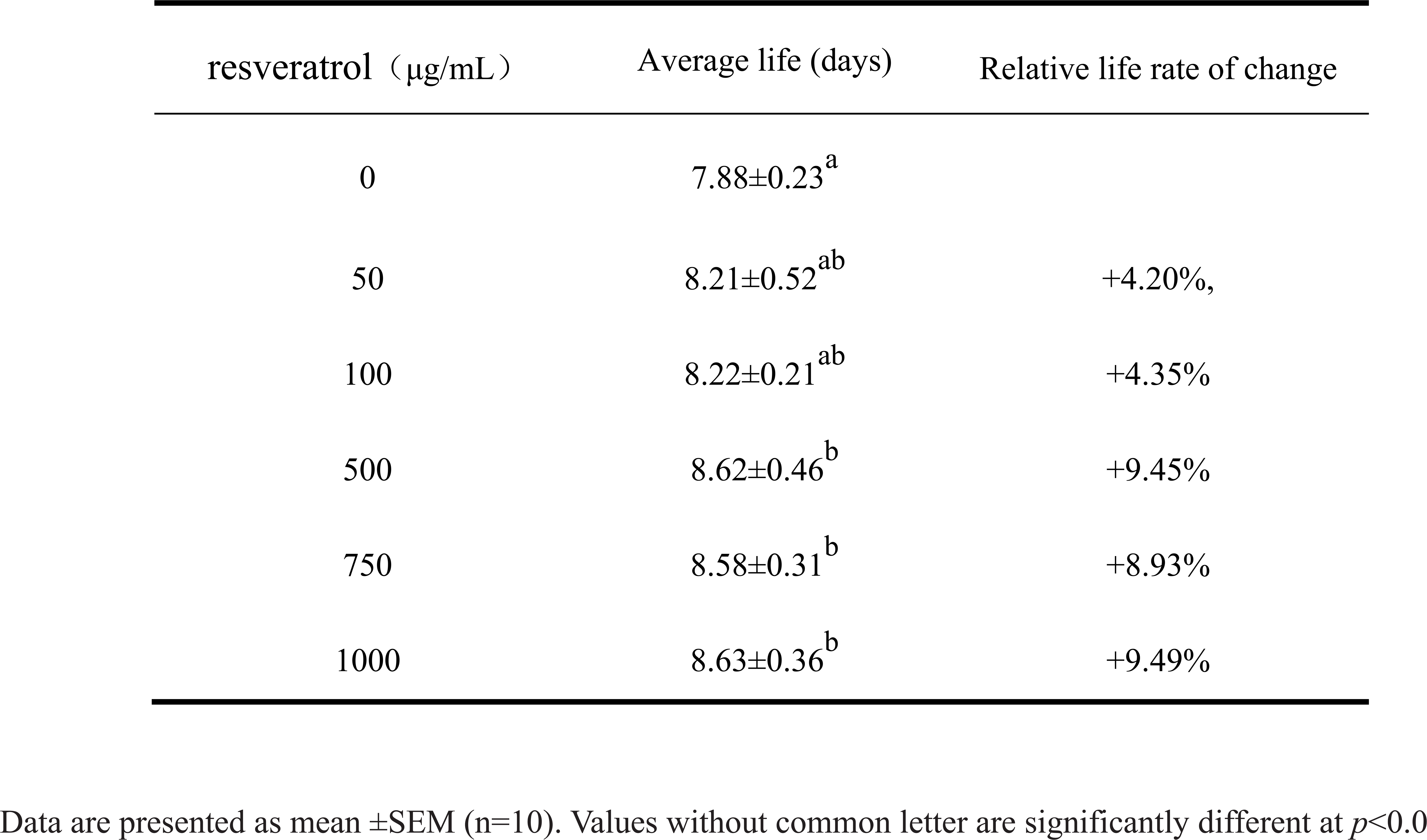
The mean life span of sugar ester damaged N2 in different concentrations of resveratrol

Similarly, we also examined the effect of resveratrol on the reproductive capacity of nematodes. The number of eggs laid by nematodes after treatment with different concentrations of resveratrol is shown in Fig.3A. It can be seen that as the concentration of resveratrol increases, the number of eggs laid by nematodes increases initially, and then, decreases. Moreover, only resveratrol treatment at a concentration of 500 mg/mL led to significant differences in the number of eggs laid compared to those in the control group.

**Figure 3.**
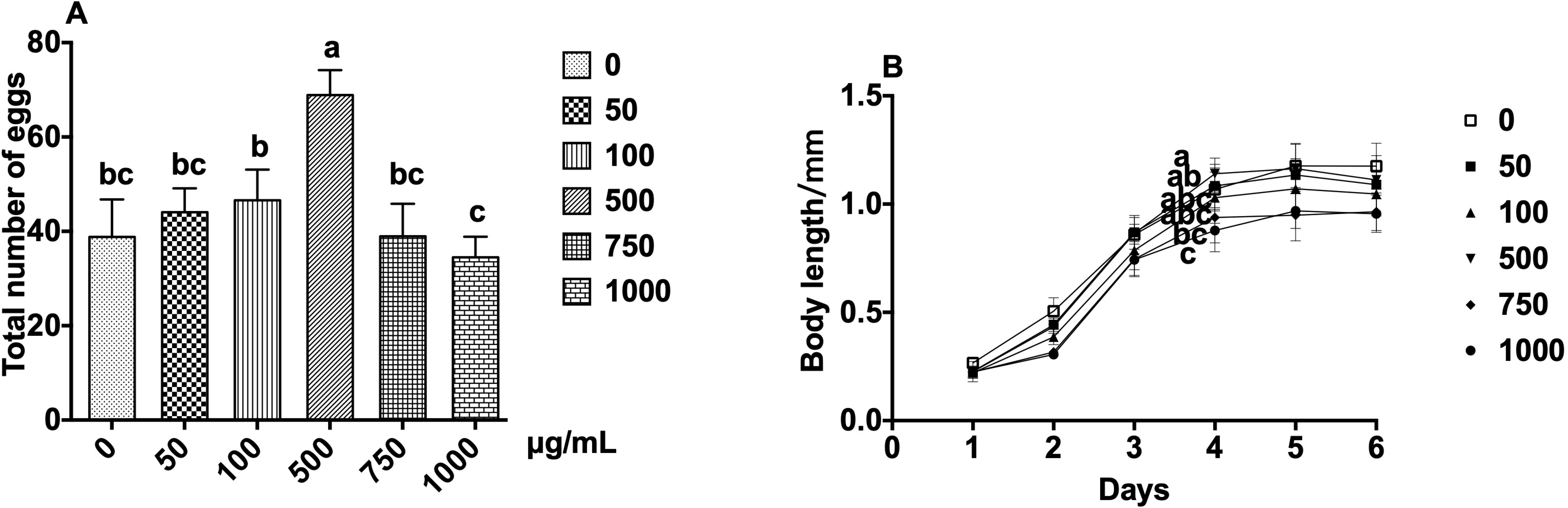
Effects of resveratrol on reproduction and body length of nematode. **A**, the effects of resveratrol on the reproductive capacity of nematodes. **B**, the effects of resveratrol on the body length of nematodes. Data are presented as mean ±SEM (n=10). Values without common letter are significantly different at p<0.05.

We determined the reparative effect of resveratrol on sucrose-stearic acid effect to shorten the length of nematodes. Contrary to what we expected, co-treatment with different concentrations of resveratrol had a synergistic inhibitory effect on the inhibition of nematode growth, which is particularly evident in the growth phase of the nematode (Fig.3B). In addition, we observed that resveratrol treatment at a concentration of 1000 µg/mL not only severely inhibited nematode development, but also significantly shortened the body length of adults.

### Differential gene expression analysis

Using Illumina sequencing technology, a survey was carried out to analyze gene expression of nematodes treated with sucrose, stearic acid, sucrose-stearic acid, sucrose-stearic-resveratrol, and control nematodes. Reads were obtained for each sample using Illumina Hiseq X Ten sequencing. After discarding the low-quality reads, corresponding to 48 million clean reads obtained from sequencing were mapped on the reference genome of *C. elegans* (GCF_000002985.6) (Table 5). High Pearson’s correlation coefficients of FPKM distribution between the three biological replicates for each sample were detected (R^2^ = 0.93-0.99, *P* < 0.001) (Figure S3), reflecting the robustness of our library preparation from nematodes’ RNA samples.

**Table 5.**
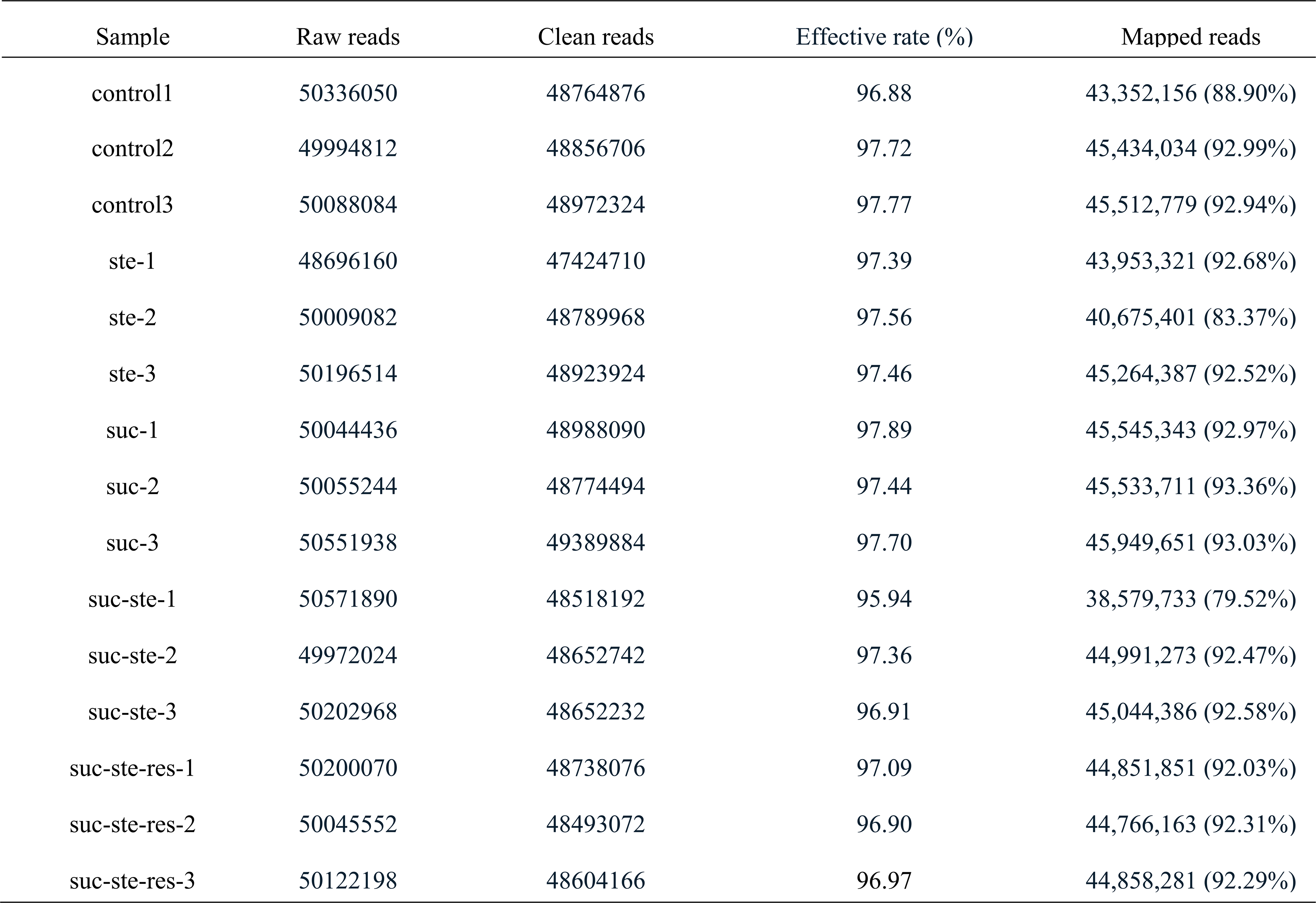
RNA-seq raw reads and alignment statistics

As shown in Table 6, compared to those in the control group, there were 905 DEGs in the sucrose group (SUC), of which 387 genes were upregulated (2-fold change, *P* < 0.05) and 518 genes were downregulated (0.5-fold change, *P* < 0.05). Similarly, there were 698 DEGs in the stearic acid group (STE), including 367 upregulated and 331 downregulated DEGs. By comparing the number of DEGs, we found that group SUC contains more DEGs than group STE, which indicated that high sucrose treatment has a more pronounced effect on nematodes than high stearic acid treatment. This is consistent with the results for the previous phenotypic indicators. Unlike in the control group, there were 1014 DEGs in group SUC-STE, including 476 upregulated DEGs, and 538 downregulated DEGs. Moreover, in contrast to the sucrose-stearic acid group, there were 10 DEGs in group REV, including 5 upregulated DEGs and 5 downregulated DEGs.

**Table 6.**
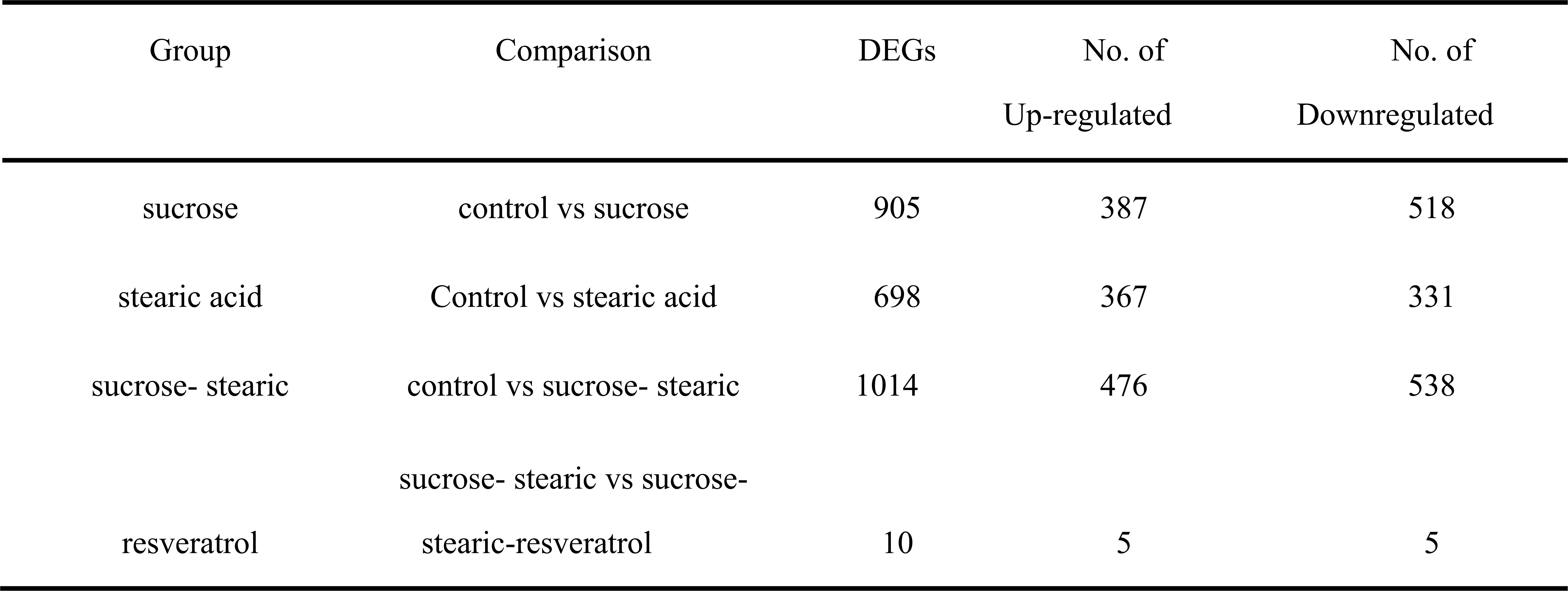
The differentially expressed genes list

### GO functional enrichment KEGG pathway analysis of DEGs

To further elucidate the gene functions, we performed GO functional analysis of the DEGs. All DEGs were assigned to three major functional categories: biological process, cellular component, and molecular function. The DEGs of comparison groups A, B, C, and D, were enriched to 27, 27, 29, and 7 subcategories, respectively (Fig.4A-D). The DEGs of comparison groups A, B, and C were mainly enriched to membrane in cellular component category, catalytic activity, and binding in molecular function category, and metabolic process, single-organism process, and cellular process in biological process. As depicted in Fig.4D, among the molecular function category, the DEGs of comparison D were more related to catalytic activity (three genes), and three genes were related to metabolic process in biological category.

**Figure 4.**
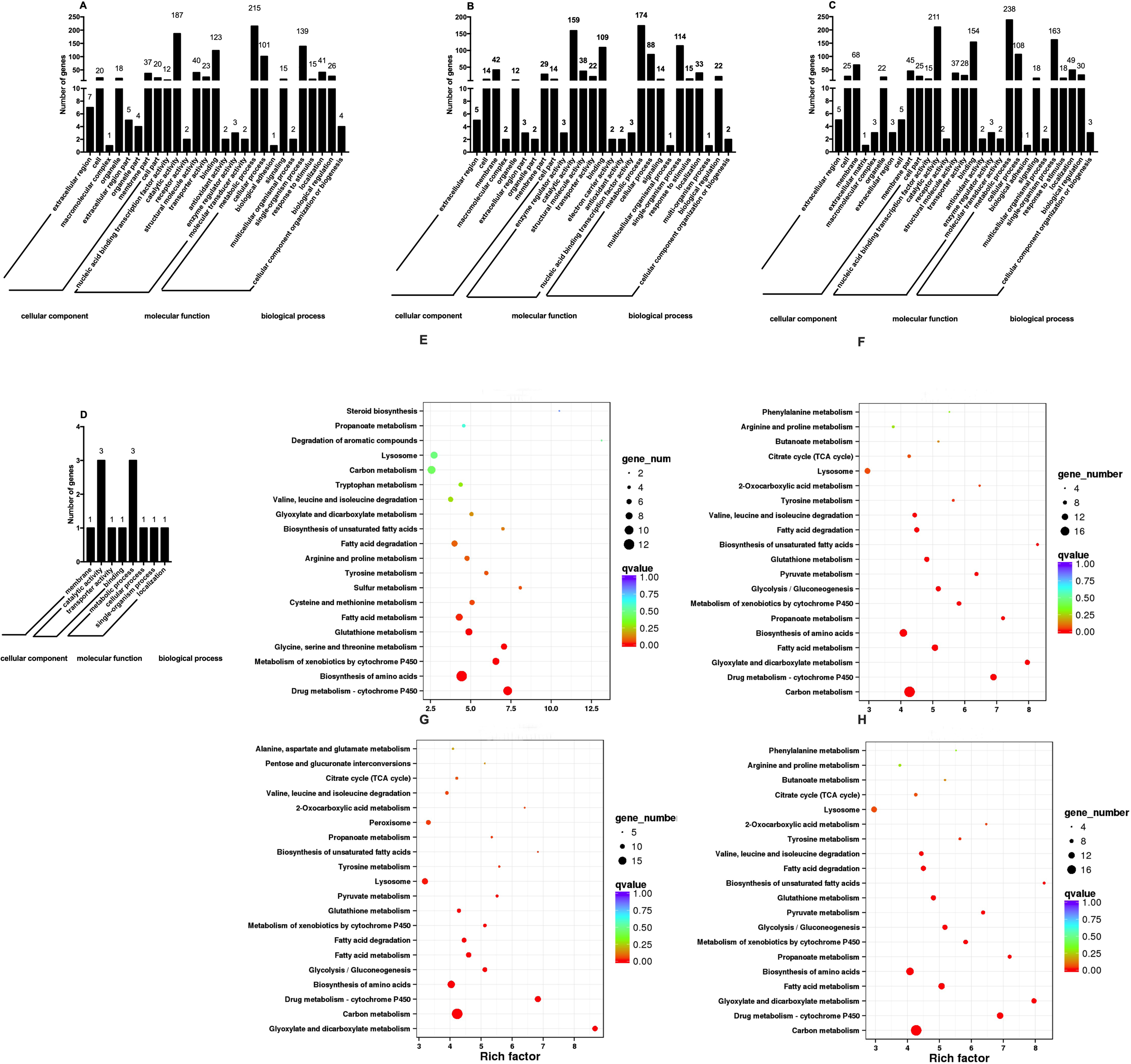
Analysis of Gene Ontology terms and KEGG pathway enrichment. Gene Ontology terms for DEGs grouped into functional categories. **A,** comparison between sucrose and control. **B**, comparison between stearic acid and control. **C**, comparison between sucrose-stearic acid and control. **D**, comparison between sucrose-stearic acid-resveratrol and sucrose-stearic acid. KEGG pathway enrichment analysis of differentially expressed genes. **E**, comparison between sucrose and control. **F**, comparison between stearic acid and control. **G**, comparison between sucrose-stearic acid and control. **H**, comparison between sucrose-stearic acid-resveratrol and sucrose-stearic acid.

We performed KEGG enrichment analysis providing in-depth insight into the biological functions of the DEGS. By using KeggArray software, all DEGS were assigned to five specific pathways, including cellular processes, environmental information processing, genetic information processing, metabolism, and organism systems. Most pathways were involved in primary metabolic processes, such as carbohydrate metabolism, amino acid metabolism, and lipid metabolism. As depicted in the Fig.4A, the genetic changes in nematodes after sucrose treatment are mainly concentrated in carbon metabolism, amino acid synthesis, and glucose metabolism signaling pathways. Stearic acid treatment led to alterations in the genes of nematodes involved in cytochrome P450-related metabolism, biosynthesis of amino acid, and fat catabolism (Fig.4B). In addition, after sucrose and stearic acid co-treatment, the genetic changes in nematodes are mainly concentrated in carbon metabolism and amino acid synthesis (Fig.4C). Interestingly, after resveratrol treatment of nematodes, only one gene (gst-25) was enriched into three metabolic pathways, including glutathione metabolism, drug metabolism, and metabolism of xenobiotics by cytochrome P450 (Fig.4D). These results further indicated that nematodes underwent active metabolic processes after treatment with sucrose and stearic acid.

## Discussion

The natural active substance resveratrol has been proved to have antioxidant, delay aging, antibacterial, anti-inflammatory, and other biological activities (Chen *et al*. 2013). However, the protective effect of resveratrol on sugar and lipid damage and its mechanism of action are still unclear. Therefore, we explored on the protective effect of resveratrol using a high sugar and high lipid model in *C. elegans*. Overall, the results indicated that low concentrations of sugars and lipids can extend nematode lifespan and promote nematode growth and development. Since, nematodes need to consume energy during growth and development, a certain amount of sugar and lipids is used to provide nutrients for nematode life activities. However, excessive sugar and lipid treatment severely shortens the life and length of the nematode and destroys its reproductive capacity. Interestingly, we found that sugar has an adverse effect on nematodes at low to medium concentrations, while lipids only cause damage to nematodes at moderate to high concentrations. In other words, sugar has a stronger effect on nematodes than lipids. Sequencing data also corroborated this result, because the differential genes generated by sugar treatment are significantly more than lipids (Table 6). In addition, the results of orthogonal tests indicated that co-treatment with high concentrations of sucrose and stearic acid had a synergistic effect on nematode damage. Nevertheless, a certain concentration of resveratrol can alleviate the damage of nematodes treated with high concentrations of sucrose and stearic acid.

Moreover, we further explored the mechanism of high sucrose and high stearic acid damage on *C. elegans* and the repair effect of resveratrol using transcriptome sequencing technology. After treatment with sucrose, stearic acid, and sucrose-stearic acid, a total of 905,698 and 1014 DEGS were identified, respectively. It suggests that high-sucrose and high-stearic acid treatment causes imbalance in nematode glycolipid metabolism by altering of the expression of several genes.

### Over-nutrition converts fat storage and exacerbates β-oxidations of fatty acids

The regulation of lipid metabolism in *C. elegans* is influenced by the environment, such as temperature and nutrient deficiencies, as well as its own physiological state, including growth, reproduction, development, and aging (Watts and Ristow 2017). Moreover, the nematode body undergoes rapid changes to produce an adaptive response to this stimulus. In our experiments, high sugar and high fat provides excess nutrients to the nematodes. In addition to digestion and absorption to meet basic life activities, nematodes store excess energy in the form of lipid droplets. In early embryos, lipid droplets are abundant, providing precursors for membrane synthesis during rapid cell division as well as energy for cellular processes until hatching. TAGs are the major component of lipid droplets, as effective energy storage molecules due to their reduced state. During fat synthesis, diacylglycerol acyltransferase encoded by dgat-2 catalyzes the formation of TAG from fatty acyl-CoA and diacylglycerols (Xu *et al*. 2012). After high-sucrose treatment and high-stearic acid treatment, nematodes convert excess nutrients into fat, by upregulating dgat-2 genes involved in TAGs synthesis. In addition, vit-2, which is involved in the transfer of dietary lipids to lipid droplets, promoting fat accumulation, was upregulated after high-sucrose treatment and high-stearic acid treatment.

Fatty acids are separated from TAGs molecules, releasing energy by β-oxidations. Fatty acids need to be activated before they enter the mitochondria for oxidation, which involves four enzymes, which are acyl-CoA dehydrogenase in the mitochondria or acyl-CoA oxidase in peroxisomes, enoyl-CoA hydratase, 3-hydroxylacyl-CoA dehydrogenase, and 3-ketoacyl-CoA thiolase (von Reuss *et al*. 2012). The genes encoding these enzymes, including acox-1, maoc-1, dhs-28, and daf-22, were upregulated in our results, which indicated that high-intensity energy metabolism was being carried out in the nematode (Figure S4).

### Fatty acids are synthesized *de novo* for growth, development, spawning, and signaling molecules

The characteristic of lipid metabolism in *C. elegans* is the synthesis of fatty acids *de novo* from acetyl-CoA. In addition to the oxidation of fatty acids to produce acetyl-CoA, other nutrients, such as carbohydrates and amino acids, can be broken down into acetyl-CoA for *de novo* fatty acid synthesis. During fatty acid synthesis, the pod-2 encoded ACC enzyme limits acetyl-CoA to malonyl-CoA transformation (Kniazeva *et al*. 2004). In the second step, the *de novo* synthesis of the fatty acyl chain by the two-carbon subunit acetyl-CoA is accomplished by the catalysis of a fatty acid synthase encoded by fasn-1 (Jia *et al*. 2016). In our study, high-sucrose treatment, high-stearic acid treatment, high-sucrose, and high-stearic acid co-treatment did not affect the expression of pod-2 and fasn-1 genes (Figure S5).

Nematodes are rich in polyunsaturated fatty acids (PUFAs), produced by desaturation. There are four fatty acid desaturases that convert 18:1n-9 into a series of C18 and C20 PUFAs, including FAT1 (Δ12), FAT2 (Δ12), FAT3 (Δ12), and FAT4 (Δ5) (Watts 2016). These polyunsaturated fatty acids provide precursors for the growth and reproduction of nematodes, and are used to synthesize fat. *C. elegans*, which is severely deficient in polyunsaturated fatty acids, exhibits many growth, reproduction, and neurological deficits. The Δ12 desaturase fat-2 mutant contained only 1% PUFAs. These mutants grow slowly, have smaller embryos, and exhibit less coordinated motion than wild-type individuals (Lesa *et al*. 2003). The Δ6 desaturase fat-3 mutant contains C18 PUFAs but does not contain C20 PUFAs. Although they grew better than the fat-2 mutant and showed a higher brood size, they showed many defects compared to the wild type (Watts *et al*. 2003). Fat-4 and fat-1 mutants contain different types of PUFAs and different proportions of omega-6 and omega-3, although their growth, development, and reproduction are largely unaffected (Watts and Browse 2002). In our experiments, high sucrose treatment and high-stearic acid treatment significantly upregulated genes encoding desaturase, including fat-1, fat-2, fat-3, fat-4, and fat-5 (Figure S5). This indicated that nematodes produce large amounts of PUFAs for growth and development through desaturation. The results of KEGG also demonstrated that high-stearic acid treatment lead to nematode lipid metabolism and decomposition disorders, affecting the growth and development of nematodes

In addition to affecting the growth and development of nematodes, PUFAs are also used as signal molecules, released from the membrane by phospholipase hydrolysis and further metabolized to form signaling molecules, collectively known as eicosanoids (Deline *et al*. 2013). In mammals, the synthesis of eicosanoids requires the participation of cyclooxygenase, lipoxygenase, and cytochrome P450 enzymes (Spector and Kim 2015). The cyp-gene family is reported to be responsible for encoding cytochrome P450s, NADPH-dependent monooxygenases that metabolize endogenous and exogenous compounds (Menzel *et al*. 2005). Sucrose treatment, stearic acid treatment and sucrose-stearic acid co-treatment of nematodes downregulated cyp-gene expression, such as cyp-29A3, cyp-14A3, and cyp-35A4 and interfered with metabolism of nematodes.

### Increased glucose metabolism shortens nematode life

Monosaccharides are directly absorbed in the body’s metabolism. The disaccharide or polysaccharide is hydrolyzed into glucose, which participates in glycolysis to provide energy to the body, or is stored as a glycogen. In mammals, glucose transport and absorption are mediated by GLUTs and insulin signaling. The fgt-1 gene is associated with nematode glucose uptake, and there have been reports that RNAi-mediated knockdown of fgt-1 extends lifespan of nematodes (Feng *et al*. 2013; Kitaoka *et al*. 2013). Previous studies indicated that inhibition of the glycolytic enzyme, glucose phosphate isomerase 1 (GPI-1), prolongs lifespan. Feng et al. (Feng *et al*. 2013) showed that disrupting glucose transport, by inhibiting FGT-1, is associated with AGE-1 and DAF-2 signaling to extend nematode lifespan. These previous studies also suggested that reduced glucose metabolism promotes longevity. In our experiments, the expression of fgt-1 and daf-2 genes was upregulated in sucrose treatment group and sucrose-stearic acid co-treatment group, while stearic acid treatment had no effect. This indicated that the addition of sucrose increased the metabolic burden of nematodes, resulting in a shortened life.

### Genes involved in the TGF-β signaling pathway

Transforming Growth Factor-β (TGF-β) superfamily ligands participate in cell identify, growth, and development. In *C. elegans*, five such ligands have been identified, including dbl-1, daf-7, unc-129, tig-2 and tig-3. Here, we only discussed dbl-1 and daf-7 signaling pathway, because their function has been explained more clearly. The core components of the dbl-1 pathway are the dbl-1 ligand, daf-4 and sma-6 receptors, and sma-2, sma-3, and sma-4 intercellular signals. Studies have shown that the lack of dbl-1 signaling pathway leads to small body size and male tail abnormal morphology (Padgett *et al*. 1998). In our gene expression profile, high-stearic acid treatment upregulated gene expression levels of dbl-1, daf-4, sma-10, and sma-6, and high-sucrose treatment slightly upregulated the expression of these genes (Figure S6), which suggested that dbl-1 signal was enhanced. Furthermore, the expression of the intercellular signals sma-2, sma-3, and sma-4 was upregulated after high-stearic acid treatment, whereas sma-2 and sma-4 were downregulated in the high-sucrose treatment group. This may be related to high-sucrose-induced shortening of nematode length. In addition, it has been reported that overexpression of the dbl-1 gene shortens the lifespan of nematodes (Luo *et al*. 2009). Both high-sucrose and high-stearic acid treatment enhanced the dbl-1 signaling pathway, which may be responsible for the shortened lifespan of nematodes caused by high sucrose and high stearic acid. daf-7, a ligand for the TGF-β signaling pathway, is involved in regulating nematodes entering the dauer phase. The core components of daf-7 pathway are: the daf-7 ligand, daf-1 and daf-4 receptors, daf-8, daf-3, and daf-14 transcription factors (Gumienny and Savage-Dunn 2013). Our results indicated that high-stearic acid treatment significantly upregulated the gene expression of daf-7, but high sucrose significantly inhibited its expression, suggesting that high sucrose may cause some stress on nematodes. There was no significant change in the expression levels of receptors daf-1 of daf-7, and there was a significant increase in daf-4 expression. Daf-8 and daf-14 act as signaling molecules, both of which are upregulated under high-stearic acid treatment and downregulated under high sucrose treatment, similar to daf-7. This indicated that high stearic acid inhibited nematodes from entering the dauer phase, while high sucrose may cause certain stress, which might promote entry of nematodes into dauer phase.

### Genes involved in the insulin signaling pathway

The *C. elegans* insulin signaling pathway (ISS) links energy metabolism with life activities, including growth, development, reproductive, longevity, and behavior (Murphy and Hu 2013). This fundamental pathway is regulated by insulin-like peptide (ILPs) ligands that bind to the insulin/IGF-1 transmembrane receptor (IGFR) ortholog daf-2. The main components of the *C. elegans* insulin signaling pathway include ILPs (Hua *et al*. 2003). Several ILPs have been shown to be involved in growth, longevity, and dauer formation of nematodes, such as daf-28 and ins gene family. In our study, we found that sucrose treatment and stearic acid treatment, sucrose-stearic acid co-treatment, and resveratrol treatment had no effect on insulin signaling pathway-related genes (daf-2, age-1, akt-1, ddl-1, hsf-1, and daf-16), except for the lipid treatment which upregulated daf-2. We only evaluated the gene expression profile of a nematode before it entered the spawning stage, and more experiments are needed to further investigate how sugar and lipids affect lifespan of nematode. In *C. elegans*, skn-1, the ortholog of Nrf-2, downstream regulator of daf-2, is required for both oxidative stress resistance and anti-aging through its accumulation in the intestinal nuclei to promote the detoxication target genes (Li *et al*. 2017). Stearic acid treatment significantly upregulated the expression of ins-27, ins-33, daf-2, and skn-1 genes. Intriguingly, sucrose treatment and sucrose-stearic acid co-treatment significantly downregulated skn-1 gene expression. This was also consistent with the phenotypic results where stearic acid was less harmful to nematode life, reproductive capacity, and body length, compared to sucrose. In addition, sugar and lipid treatments downregulated genes (gst gene family and ugt gene family) related to oxidative stress. Furthermore, in our gene expression profile, the acdh-1 gene encoding the short-chain acyl-CoA dehydrogenase in mitochondria was upregulated after high glucose and high fat treatment. This may result in increased mitochondrial activity, increased rate of oxidative phosphorylation, increased metabolism, and reduced lifespan.

### Resveratrol protects sugar and lipid damage to nematodes

UDP-glycosyltransferase catalyzes the transfer of glycosyl groups from activated donor molecules to receptor molecules, and participates in several activities, such as detoxification, defense response, and regulation of hormone levels (Zhou *et al*. 2018). Glutathione S-transferase reduces cellular oxidative stress. Comparing differential gene analysis of sucrose-stearic acid co-treatment group and resveratrol group, we found that the repair effect of resveratrol on damaged caused by sucrose-stearic acid treatment on nematodes may be related to UDP-glycosyltransferase and glutathione S-transferase. KEGG analysis showed that the repair of resveratrol may be related to the metabolism of cytochrome P450 to foreign substances and glutathione metabolism (Fig.4D). Our results were consistent with previous studies which reported that resveratrol acts against oxidative stress by regulating cytochromes involved in the metabolism of exogenous substances (Fischer *et al*. 2017). Taken together, we speculated that the repair effect of resveratrol on damage due to high sucrose-stearic acid is mainly manifested in two aspects; one is to reduce the oxidative stress of cells and the other is to participate in the metabolism of exogenous substances.

## Conclusion

Intake of a certain amount of sugar and lipid promotes the growth and development of nematodes and prolongs their life to some extent. However, excess sugar and lipid intake disrupts the metabolism of nematodes, causing a certain degree of damage to their longevity, growth, and reproduction. Moreover, the high sugar phase causes more severe damaged than the high lipid phase, mainly due to an increase in the metabolic burden of nematodes and interference with normal metabolic function. The protective effect of resveratrol on nematodes is manifested as follows: reduction of cellular oxidative stress and participation in the metabolism of exogenous substances. Resveratrol is expected to be used to alleviate damage to the body due to over-nutrition.

## Declarations of interest

There were no conflicts of interest in this study.

## Acknowledgements

This study was supported by National Natural Science Foundation of China (Grant No. 31790412)

## Authors’ contribution

Funding acquisition, Yali zhang; Methodology, Lin Zhang; Resources, Xiong Wang, Lin Zhang, Lei Zhang, Sihan Wei, Wenli Wang and Jie Wang; Supervision, Huilian Che and Yali zhang; Writing – original draft, Xiong Wang; Writing – review & editing, Xiong Wang.

## Supplementary materials

**Figure S1**. The effects of sucrose(A), fructose (B) and glucose (C) on the lifespan of nematodes. Values without common letter are significantly different at *p*<0.05.

**Figure S2**. The effects of stearic acid (A), linoleic acid (B) and cholesterol (C) on the lifespan of nematodes. Values without common letter are significantly different at p<0.05.

**Figure S3**. Analysis of sample expression correlation after transcriptome sequencing

**Figure S4**. FPKM value of different genes related to fat storage and exacerbates β-oxidations of fatty acids. Values without common letter are significantly different at *p*<0.05.

**Figure S5**. FPKM value of different genes related to fatty acids synthesized. Values without common letter are significantly different at *p*<0.05.

**Figure S6**. FPKM value of different genes involved in the DBL-1 signaling pathway. Values without common letter are significantly different at *p*<0.05.

**Figure S7.** FPKM value of different genes involved in the DAF-7 signaling pathway. Values without common letter are significantly different at *p*<0.05.

**File S1.** Relative expression of genes (GSTs, CLECs, COLs, CYPs, GRLs, LYSs) in each treatment.

